# Anillin Related Mid1 as an Adaptive and Multimodal Contractile Ring Anchoring Protein: A Simulation Study

**DOI:** 10.1101/2023.01.27.525865

**Authors:** Aaron R. Hall, Yeol Kyo Choi, Wonpil Im, Dimitrios Vavylonis

## Abstract

The organization of the cytokinetic ring at the cell equator of dividing animal and fungi cells depends crucially on the anillin scaffold proteins. In fission yeast, anillin related Mid1 binds to the plasma membrane and helps anchor and organize a medial broad band of cytokinetic nodes, which are the precursors of the contractile ring. Similar to other anillins, Mid1 contains a C terminal globular domain with two potential regions for membrane binding, the Pleckstrin Homology (PH) and C2 domains, and an N terminal intrinsically disordered region that is strongly regulated by phosphorylation. Previous studies have shown that both PH and C2 domains can associate with the membrane, preferring phosphatidylinositol-(4,5)-bisphosphate (PIP_2_) lipids. However, it is unclear if they can simultaneously bind to the membrane in a way that allows dimerization or oligomerization of Mid1, and if one domain plays a dominant role. To elucidate Mid1’s membrane binding mechanism, we used the available structural information of the C terminal region of Mid1 in all-atom molecular dynamics (MD) near a membrane with a lipid composition based on experimental measurements (including PIP2 lipids). The disordered L3 loop of C2, as well as the PH domain, separately bind the membrane through charged lipid contacts. In simulations with the full C terminal region started away from the membrane, Mid1 binds through the L3 loop and is stabilized in a vertical orientation with the PH domain away from the membrane. However, a configuration with both C2 and PH initially bound to the membrane remains associated with the membrane. These multiple modes of binding may reflect Mid1’s multiple interactions with membranes and other node proteins, and ability to sustain mechanical forces.

## Introduction

Cytokinesis, the last step in a cell cycle, is a fundamental biological process in animals and fungi that describes the division of a mother cell into two daughter cells, aided by the formation and contraction of an actomyosin ring. In animal cells, the highly conserved protein anillin is involved in the coordination and localization of cytokinetic ring components and, additionally, anchors the ring to the plasma membrane [1–3]. While the N-terminus of anillin is known to bind ring components, its C-terminal region associates to the plasma membrane, through PIP_2_ contacts in particular [4]. The C-terminal region of anillin is comprised of a C2 domain, PH domain, and Rho binding domain (Supplemental Figure S1A, B). The C2 domain contains a nuclear localization sequence (NLS), mutation of which reduces anillin’s affinity for the equatorial cortex [5, 6].

In model organism *Schizosaccharomyces pombe* (fission yeast), ring positioning is driven by anillin related protein Mid1 [7], whose spatial localization in the cell middle is regulated by positive cues, such as nuclear shuttling [8] and affinity for PIP_2_ that is enriched in the cell middle during mitosis [9], and negative cues, such as phosphoregulation by the kinase Pom1 that localizes at the cell poles away from the cell middle [10]. At the cell center Mid1 organizes cytokinetic “nodes,” primarily consisting of myosin Myo2, formin Cdc12, F-BAR domain containing Cdc15, and IQGAP Rng2 [11]. Mid1 is an important upstream component that aids in the recruitment of Myo2 and Cdc15 [12]. After a period of node maturation, the medial band of nodes condenses into the cytokinetic ring through Cdc12-driven actin filament elongation and Myo2-driven contraction [13]. Eventually Mid1 leaves the ring prior to its constriction [14]. Membrane connections for both Cdc12 and Myo2 are important in the transmission of force for node movement, although the specifics of the force generation is poorly understood [15]. While Mid1 deletion is not lethal, *mid1Δ* cells form misplaced and tilted cytokinetic rings, which are otherwise tightly regulated to form at the cell center and perpendicular to the long axis of the cell [16–19]. Additionally, cells lacking Mid1 divide slower and at a more variable rate, resulting in cells of more variable length and a higher incidence of multinucleation [20]. Furthermore, colocalization of important ring components is disrupted and the cytokinetic ring appears to form in a mechanistically dissimilar way to wildtype cells [19]. Super-resolution studies indicate that Mid1 localizes closer to the membrane than any other node or ring component and therefore understanding the geometry of its membrane binding is critical to understanding the early organization of cytokinetic node and ring components [21, 22].

Structurally similar to anillin, Mid1 has a C terminal globular region that includes a C2 domain (residues 580-787, Figure 1A, B with components in cyan, dark blue, and green) connected to a PH domain (805-900, Figure 1A, B in yellow and orange). While Mid1 lacks the Rho binding domain present in anillin, the geometrical relationship between the PH and C2 domains as studied by crystallography (for Mid1 [9]) and as predicted by AlphaFold [23, 24] are similar (compare Supplemental Figure S1A and B to C and D and Figure 1A). In Mid1, the PH and C2 domains are connected by an intermediate region referred to as a connector (CNCT) domain (787-804, 901-920, Figure 1A, B in red) not present in anillin. The Mid1 C2 domain includes a notably long L3 loop (654-710, Figure 1A, B in dark blue) with a highly negatively charged NLS (691-696, Figure 1A, inset), similar to anillin, near a predicted amphipathic helical structure [23–25] (“candidate helix”, 681-688, Figure 1A, inset), which may localize to the plasma membrane. The structure of the majority of the PH, C2, and CNCT domains (Mid1 C2-PH) has been resolved through X-ray crystallography, excluding the L3 loop and the smaller ß7-ß8 loop (“L0 loop”, 745-760, Figure 1A, B in green). In contrast to anillin, Mid1 dimerizes through an interface on its C2 domain, and the structure of a Mid1 C2-PH dimer has been resolved crystallographically (Supplemental Figure S2A, B) [9]. A long Mid1 N-terminal region, predicated intrinsically disordered (1-579, Figure 1A in grey, not depicted in B), has been shown to be functionally interchangeable with human anillin’s N-terminal region [9].

**Figure 1:**
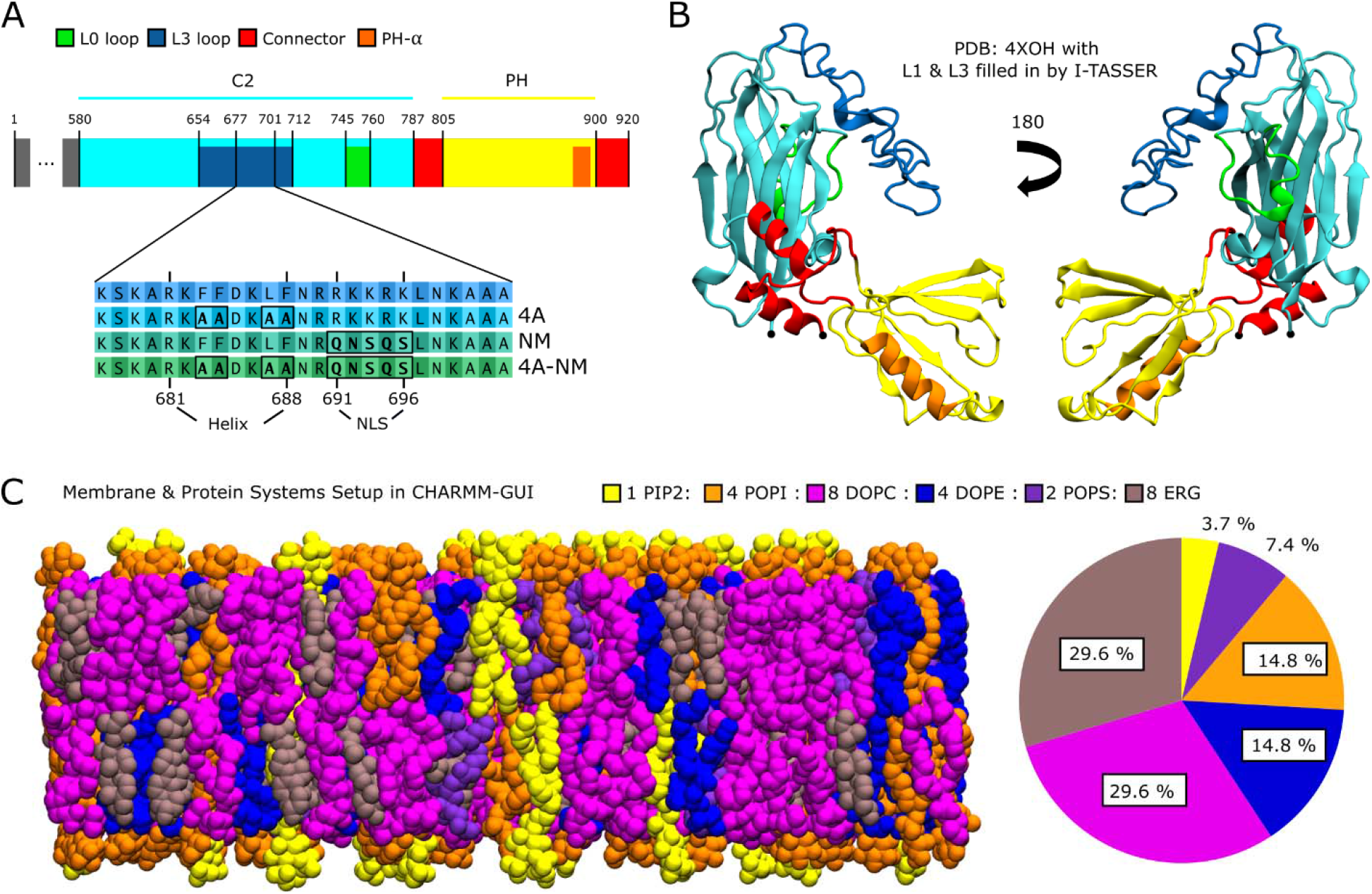
Mid1 Protein and Membrane System. **(A)** Mid1 sequence. L3 mutations investigated defined in sequence blowup. **(B)** Mid1 structure from PDB: 4XOH [9]. Missing residues that make up the L0 & L3 loop were filled in using I-TASSER [35]. Visualization of protein and membrane systems done in VMD [62]. **(C)** Example snapshot of membrane bilayer with composition used in all simulations (left). Legend lists the molar ratio of membra**ne** constituents; also depicted pictorial (right), selected to match [36]. Tail saturation was independently matched to [37]. Protein and membrane systems were prepared using CHARMM-GUI (see Methods).

Both the Mid1 C2 and PH domains have been indicated in membrane binding. Purified Mid1 PH have been shown to bind to negatively charged phosphatidylinositol (PI) lipids, including PIP2 that is present in higher concentration towards the medial region of fission yeast [26]. When the PH domain is truncated, Mid1 cortical localization is disrupted in cells lacking Cdr2, a component of medial interface nodes that precede cytokinetic nodes [26]. In addition, mutations to the L3 loop’s NLS sequence and candidate helix have striking effects on the localization of Mid1 during the cell cycle [25]. Notably, simulation studies have described the binding of PH and C2 domains of various other proteins to PI lipids [27–31].

The orientation of Mid1 when binding the plasma membrane as well as its stoichiometry as a monomer or as a dimer remain unresolved. In the Mid1 C2-PH dimer crystal structure, the L3 loops, one from each C2 domain, are near each other and point in the same direction; in contrast, the PH domains are distant from each other and orient in different directions (Supplemental Figure S2A). Given the geometry of the dimer in the crystal structure, it is difficult to imagine how the PH domains would bind to the membrane cooperatively with each other or the L3 loops, leading to the hypothesis that the Mid1 dimer membrane binding mode only involves the C2 domain/L3 loops [9]. This is consistent with the expected vertical orientation of type II C2 regions, to which Mid1’s C2 region belongs [29]. However, Mid1 could possibly bind the membrane as either a monomer or as a dimer. Indeed, an orientation for a monomer binding mode using both PH and C2 domains has been hypothesized for both human anillin [9], which is not expected to dimerize [3, 32, 33], and Mid1 [34].

To elucidate how Mid1 binds the membrane, we performed all-atom MD simulations of sections of the Mid1 monomer utilizing available structural data, with an experimentally informed membrane representation of the fission yeast plasma membrane near the cell center. Looking at results from Mid1 subdomains in isolation and together, while making comparisons to simulations of known membrane binding defective mutations, provided us with a clearer picture of how the Mid1 monomer utilizes its various subdomains to make a connection to the fission yeast cortex. These results can further inform our understanding of the Mid1 dimer’s role in anchoring the rest of the cytokinetic node to the plasma membrane, as well as more broadly to the function of anillins in cytokinesis.

## Results

In order to investigate how the Mid1 protein binds to the membrane, we performed all-atom MD simulations using the crystal structure of the C2-PH domain (PDB: 4X0H)[9] with the missing residues filled-in by I-TASSER [35] (Figure 1B). Recent studies have characterized the lipid and ergosterol composition of budding yeast [36], a closely related organism to fission yeast, and the saturation levels of lipid tails in fission yeast membrane [37]. We used a lipid bilayer composition that closely matches this data, by independently setting the molar ratio of lipid heads and ergosterol to match the ratio measured in [36] and selecting tail saturation levels to match the molar ratio in [37]. The resulting bilayer includes palmitoyl-oleoyl-phosphatidylinositol-(4,5)-bisphosphate (PIP_2_), palmitoyl-oleoyl-phosphatidylinositol (POPI), di-oleoyl-phosphatidylethanolamine (DOPE), di-oleoyl-phosphatidylcholine (DOPC), palmitoyl-oleoyl-phosphatidylserine (POPS) and Ergosterol (ERG) with molar ratios described in Figure 1C. The upper and lower leaflet composition are identical, and due to periodic boundaries, molecules can bind freely to either leaflet.

### Mid1 L3 loop and PH domain can independently bind to the membrane

We began by examining if the candidate membrane binding domains can bind a lipid bilayer with a composition similar to that of fission yeast in isolation. We performed all-atom MD of a segment of the L3 loop (residues 677-701 in Figure 1A, please see Table 1 for each system information) as this section includes both the NLS sequence and the sequence expected to bind the membrane as an amphipathic helix (from now on referred to simply as the “candidate helix”). This L3 segment is expected to be flexible and important for membrane binding [9]. Additionally, the relatively small size of the system may allow us to investigate the emergence of helix formation without interference from neighboring sections of the L3 loop. We initialized the L3 segment far enough apart from the membrane bilayer to allow it to diffuse before binding (Supplemental Figure S3A).

**Table 1:** Summary of simulation parameters, to within small fluctuations among replicas.

| System Name | System Size<br>(Å) | Number of<br>Total Atoms | Number of<br>Replicas | Total Simulation Time<br>of All Replicas (μs) |
| --- | --- | --- | --- | --- |
| L3 segment | 89×89×159 | 128,000 | 10 | 5.0 |
| L3 segment 4A | 89×89×159 | 128,000 | 10 | 5.0 |
| L3 segment NM | 89×89×159 | 128,000 | 10 | 5.0 |
| L3 segment 4A-NM | 89×89×159 | 128,000 | 10 | 5.0 |
| PH domain | 89×89×159 | 128,000 | 10 | 5.0 |
| C2-PH unbound | 145×145×200 | 418,000 | 10 | 14.8 |
| C2-PH unbound 4A | 145×145×200 | 418,000 | 2 | 3.0 |
| C2-PH unbound NM | 145×145×200 | 418,000 | 2 | 3.0 |
| C2-PH unbound 4A-NM | 145×145×200 | 418,000 | 2 | 3.0 |
| C2-PH bound | 138×138×143 | 281,000 | 10 | 3.3 |

In all ten independent simulations we performed for 500 ns, the L3 segment bound the membrane quickly (<22 ns, Movie 1). The binding of the L3 segment occurred primarily through charged lipid contacts with PIP_2_ & POPI (Figure 2A, bottom). Some contacts between the L3 loop and lipid tails were seen both in the N-terminal region and NLS sequence. However, membrane insertion near the N-terminus is likely an artifact of the truncated segment and is far less present in C2-PH simulations (see next section, Figure 3C). α-helix formation was seen in most of the predicted residues (681-688), although this occurred with less than 25% frequency. Residue K682 appears to break up helical regions, but this may be due to unphysical N-terminal membrane insertion (Figure 2A, top). However, we did not observe the expected amphipathic helix binding mode: the hydrophobic residues (F683, F684, L687, F688) in the candidate helix make very few lipid tail contacts and face mostly away from the membrane (Figure 2B). It is possible more simulation time or simulating a more precise section of the protein is required to see helix formation and membrane insertion of the hydrophobic residues in the same simulation.

**Figure 2:**
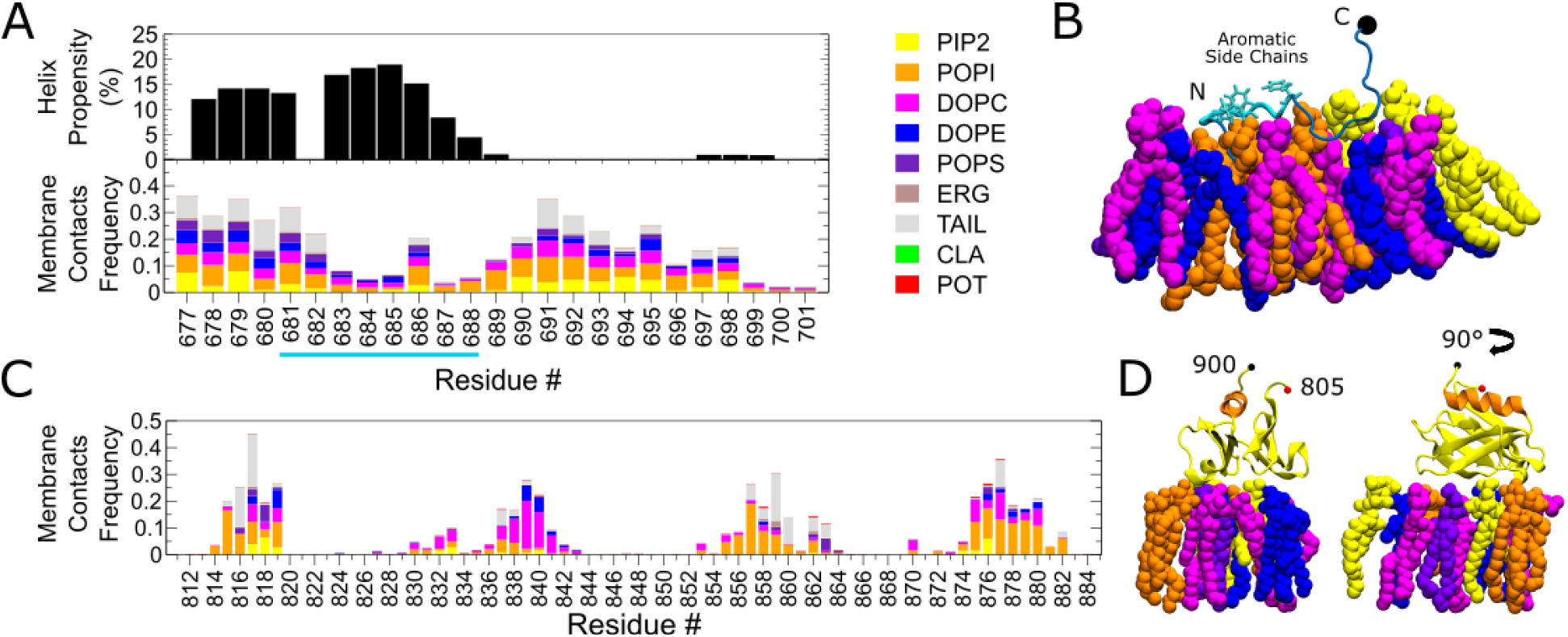
L3 loop & PH domain can independently bind to the membrane. **(A)** Ten 500 ns independent simulations of the L3 segment (residues 677-701) started apart from a lipid bilayer with composition described in Fig. 1C were performed (see Methods). The first 100 ns of each simulation were not included in the analysis. (Top) Helix propensity calculated by DSSP [64]. Residues underlined in cyan are predicted to be helical by AlphaFold [23, 24]. (Bottom) Contact frequency of L3 segment residues with membrane components. Contact frequency is normalized to 1, with the remaining contacts being water (not shown). **(B)** Simulation snapshot of L3 segment binding the membrane. Only showing membrane components within 5 Å. Membrane components are colored according to the legend in (A). Residues in cyan correspond to underlined residues in (A). **(C)** Ten 500 ns independent simulations were performed for the PH domain (residues 805-900) started apart from a lipid bilayer with composition described in Fig. 1C (see Methods). However, eight simulations featured unphysical membrane binding of terminal ends. Contact frequency of PH domain residues with membrane components were calculated using bound frames of the remaining two simulations. Contact frequency is normalized to 1, with the remaining contacts being water (not shown). **(D)** Snapshot of simulation of PH domain binding the membrane. Only showing membrane components within 5 Å. Membrane components are colored according to the legend in (A).

**Figure 3:**
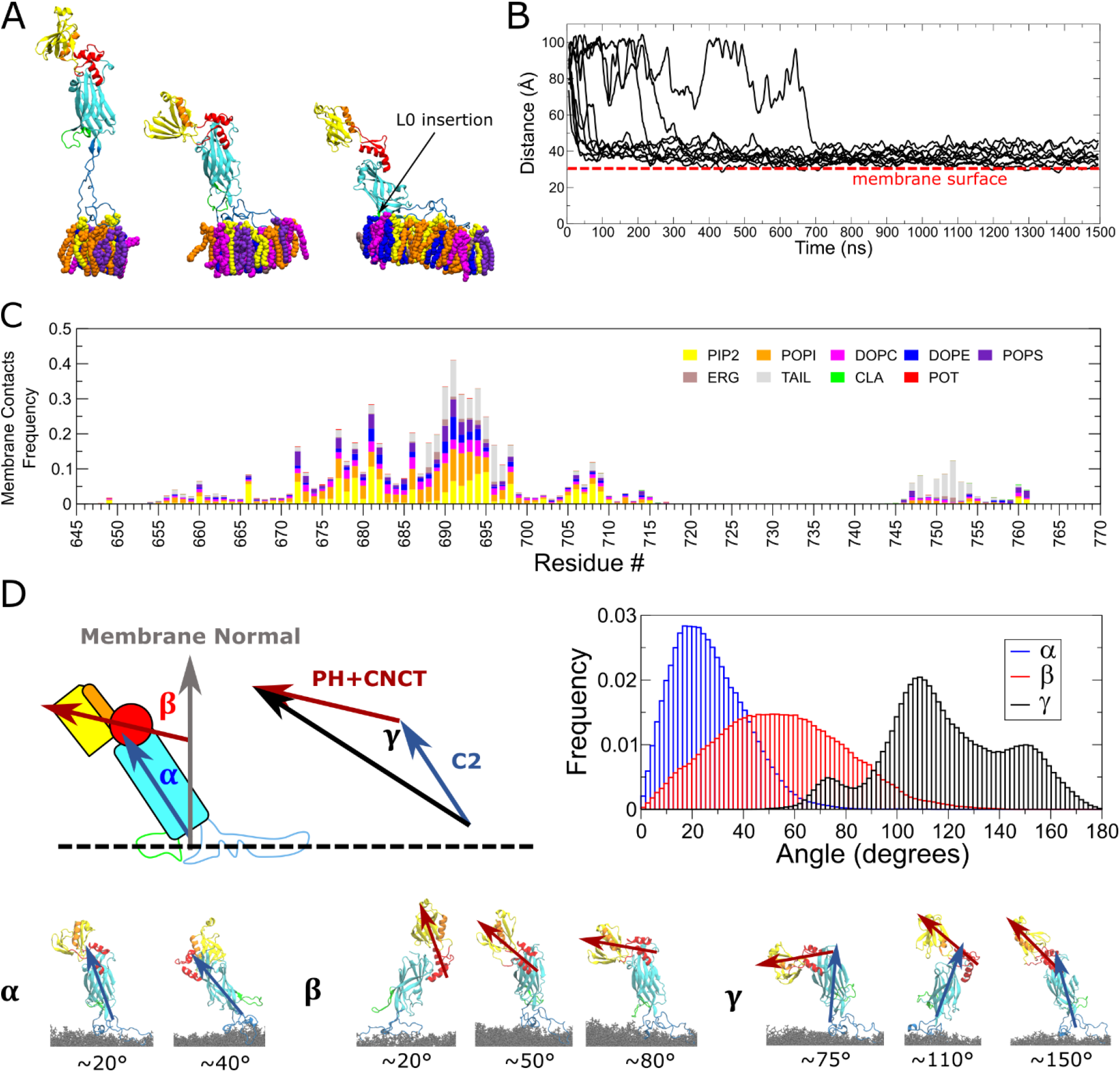
C2-PH binds the membrane quickly through the L3 loop and remains in a configuration where the PH is distal to the membrane. **(A)** Simulation snapshots of Mid1 C2-PH (residues 580-920) binding to the membrane in 3 configurations: through the L3 far from the membrane, through the L3 near to the membrane, and through the L0 & L3. Only showing membrane components within 5 Å. Membrane components are colored according to the legend in (C). **(B)** Time sequence of the distance between the L3 and membrane COM along the membrane normal. Data shown is a running average over a 100 ns window. Ten independent simulations were run for at least 1300 ns with the Mid1 C2-PH started apart from a lipid bilayer of composition described in Fig. 1C (see methods). **(C)** Contact frequency of C2-PH residues with membrane components. Analysis includes the last 1000 ns of the ten simulations in **(B).** Contact frequency is normalized to 1, with the remaining contacts being water (not shown). **(D)** Angle frequency (upper right) of defined angles α, β, and γ(upper left) in last 1000 ns of simulations in (B). Data is normalized and binned in a fixed 100 bins. α is defined as the angle between the first principal component (PC1) of the C2 domain and the membrane normal. β is defined as the angle between PC1 of the PH and connector domain (PH+CNCT) and the membrane normal. γ is defined as the inner angle of a triangle formed by PC1 of C2 and PC1 of PH+CNCT. Simulation snapshots depict representative configurations at the stated angles (bottom).

Next, we performed ten independent simulations of the PH domain (residues 805-900, Table 1) initialized far enough apart from the membrane to allow it to freely diffuse (Supplemental Figure S3B). In all ten simulations, the PH domain was able to quickly bind the membrane (<60 ns, Movie 2). However, in eight of the ten simulations, the PH domain bound and remained in contact with terminal residues, which would not be available for binding in the full-length protein. Therefore, we restricted our analysis to PH membrane binding in the two simulations which showed binding independent of the terminal regions (Figure 2C). Similar to the L3 segment, the PH domain makes many contacts with charged POPI & PIP_2_ lipids. Although it appears to form less extensive PIP_2_ contacts, this may be due to insufficient sampling. The binding conformation of the PH domain, with the PH-α distal to the membrane, is similar to other simulations studies of PH domains with the majority of contacts between PIs and the ß1-ß2, ß3-ß4, ß5-ß6, and ß6-ß7 loops [27, 29].

### C2-PH binds the membrane quickly through the L3 loop and remains in a vertical configuration with the PH is distal to the membrane

Having established that both the PH and C2’s L3 loop can bind the membrane, and characterized their respective features, we moved towards investigating if and how the C2-PH will bind when allowed to freely diffuse to the membrane. We created ten initial conditions of the C2-PH started away from the membrane to allow reorientation before binding (Supplemental Figure S3C). Each simulation had a different rotational orientation, sampled from an approximately uniformly spherical distribution, to study the mode of binding resulting from varying initial conditions, and was run for at least 1300 ns (Table 1).

In all ten simulations, the C2-PH binds initially through the L3 loop (Figure 3A, left, Movie 3). The L3 loop was continuously associated with the membrane in all cases. The C2 domain remained in a vertical orientation in all cases, though in one simulation, the C2 domain tilted in a way that allowed the L0 loop to make additional contacts and insert into membrane (Figure 3A, third snapshot, Movie 4). In this simulation the L0 binding began after 200 ns and persisted until the end of the simulation (1300 ns). Other simulations lacked L0 contacts or only showed contacts that remained at the surface of the membrane (Movie 3, at 250 ns). The majority of simulations bound in less than 100 ns (Figure 3B, Movie 3, 4).

We analyzed membrane contacts, averaged over the ten copies and using the last 1000 ns for all but one copy which we used the last 700 ns, a period over which all copies were bound to the membrane (Figure 3C). The results look nearly identical when only the last 500 ns were used for this analysis (data not shown). L3 binding primarily occurs through interactions with charged POPI and PIP_2_ lipids, especially in the NLS region. However, the candidate helix 681-688 makes notably less contacts than surrounding residues. The contacts for residues 677-701 are mostly in agreement with the previous simulations of this sequence in isolation (compare Figure 3C to Figure 2A bottom). However, residues 677-682 now make minimal tail contacts in the presence of the whole C2-PH. In general membrane insertion of the L3 loop is minimal as quantified by presence of tail contacts. A region of the L3 loop not included in the L3 segment simulations (Figure 2), 705-710, also makes strong contacts with charged POPI and PIP_2_ lipids, likely due to three positively charged lysines (K706, K708, K709). The L0 loop contacts mostly involve lipid tails, likely due to its high number of hydrophobic residues (F747, L748, A750, I751, V753, I755, I758).

The PH domain did not contact the membrane in any simulations and remained distant to the membrane during L3 binding. The C2 domain explores a range of angles with respect to the membrane normal, but this range is not large enough to bring the PH within binding distance to the membrane (Figure 3D). The connector region exhibits noticeable flexibility, which allows the PH domain to explore a wide range of angles with respect to the C2 domain (Figure 3D). However, these fluctuations were also not large enough to bring PH close to the membrane while the L3 loop is bound. Cases with L0 loop binding may cause some tilt in the C2 domain’s binding angle, but this was not enough to allow the PH domain to reach the membrane either (Figure 3D). One explanation for why the PH domain does not bind the membrane in C2-PH unbound simulations could be due to kinetics and geometry: the L3 loop’s flexibility may allow it to reach and bind the membrane more quickly than the PH domain, placing the C2-PH region in a vertical orientation with PH far from the membrane.

The results of Figure 3 reveal a new membrane binding interface of Mid1 in the L0 loop and also show that while the L3 and PH can bind readily in isolation, the context of the C2-PH provides geometric and kinetic obstacles for the PH domain. In the following sections we attempt to resolve the remaining questions regarding the lack of PH binding and the lack of strong contacts or membrane insertion in the candidate helix, which were expected from previous experimental studies [25].

### Simultaneous C2-PH domain binding is stable

From the results of the freely diffusing C2-PH simulations, we hypothesized that there may be geometric and kinetic barriers that prevent PH binding during the available simulation time either when the C2-PH is away from the membrane, or following L3 binding. In order to investigate if PH binding might occur over longer times, we initialized the C2-PH such that the L3, L0, and PH domains were in contact with the membrane in a configuration similar to that proposed by Chatterjee and Pollard [34] (Figure 4A, left). We performed ten independent simulations of at least 300 ns starting from the C2-PH bound configuration and then analyzed the data after the first 100 ns (Table 1). In all ten simulations the PH domain remained bound for the entire duration and showed no signs of detaching (Figure 4A, right, Movie 5). The L3, L0 and PH domain formed robust contacts (Figure 4B). The PH domain makes many contacts between lipids, especially POPI, and the ß1-ß2, ß3-ß4, ß5-ß6, and ß6-ß7 loops. The PH-α remains distal to the membrane as it was initialized. The PH domain contacts seen agree with those when simulated in isolation, including less prevalent contacts with PIP_2_ in comparison to the L3 loop (Compare Figure 4B with Figure 2C).

**Figure 4:**
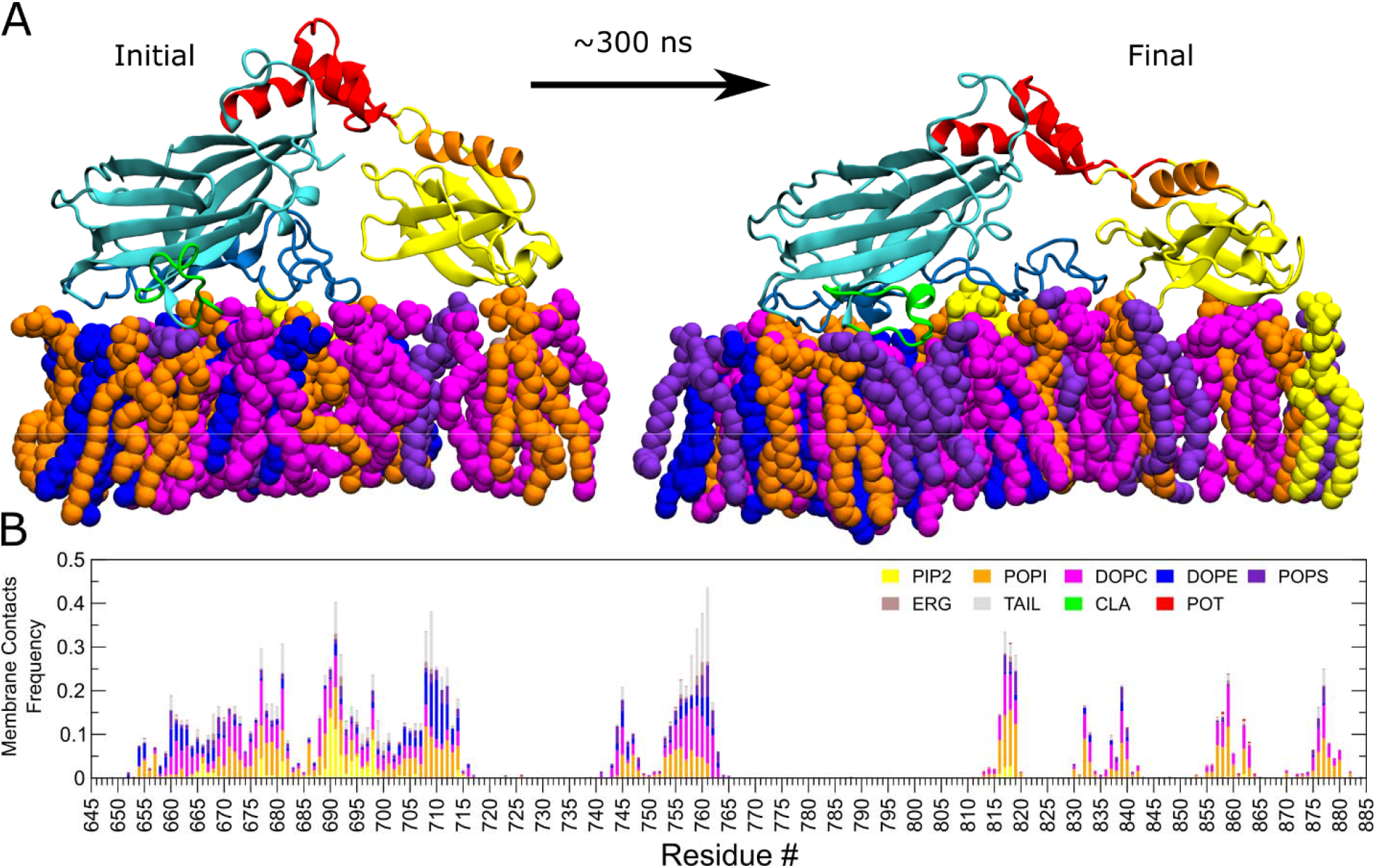
Simultaneous C2-PH domain binding is stable. **(A)** The Mid1 C2-PH (residues 580-920) was initialized near the membrane in a configuration similar to that proposed in [34]. After equilibration, 10 independent simulations were performed for at least 300 ns (see Methods). Snapshots show the initial and final configuration of one simulation. Only showing membrane components within 5 Å. Membrane components are colored according to the legend in (B). **(B)** Contact frequency of the ten independent simulations. The first 100 ns from each simulation were ignored. Contact frequency is normalized to 1, with the remaining contacts being water (not shown).

Taken together with the results of the freely diffusing C2-PH binding, these results indicate that there is a kinetic barrier preventing the PH domain from binding on short timescales, but stable simultaneous binding of C2 and PH domains might occur over long periods. Therefore, the Mid1 C2-PH has several binding interfaces it can utilize to bind stably to the membrane.

### L3 mutants can still bind membrane, but NM mutation is weaker

As mutations of the L3 loop have been shown to disrupt Mid1 cortical localization in cells [25], we were interested in how these mutations would affect membrane binding and contacts. We performed ten simulations each of the L3 segment for a duration of 500 ns under mutations to the candidate helix, in which four residues are replaced by alanines (4A, Movie 6), mutations to the NLS sequence, in which five positively charged residues are replaced with the sequence QNSQS (NLS mutant, NM, Movie 7), and a double mutant of these mutations (4A-NM, Figure 1A, Movie 8), as in the study of [25] (Table 1). The first 100 ns were excluded from the analysis by which time all simulations were bound. None of the mutations resulted in completely preventing membrane binding of the L3 segment in any of the simulations. The helix propensity of the candidate helix 681-688 remained similarly weak to WT for the 4A mutation and was even lower for the NM and 4A-NM mutations (Figure 5A). The membrane contacts of the 4A mutant are not qualitatively distinguishable from WT. However, the NM and 4A-NM mutants have noticeably less contacts with POPI and PIP_2_ in the NLS region (compare Figure 5B left with Figure 2A).

**Figure 5:**
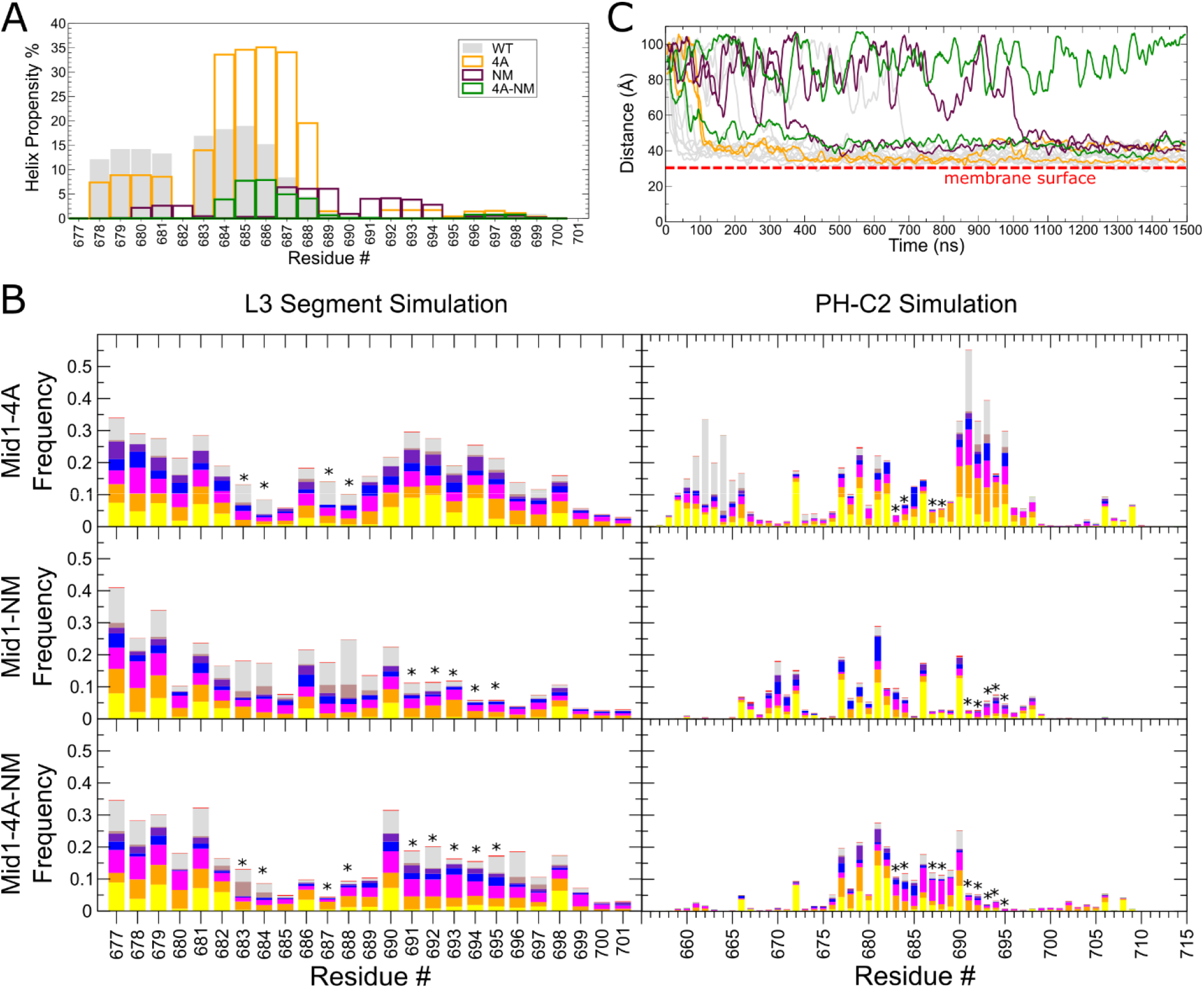
L3 mutants can still bind membrane, but NLS mutation is weaker. **(A)** Ten 500 ns simulations were performed of the L3 segment (residues 677-701) for each mutation defined in Fig. 1A. The first 100 ns of each simulation were not included in analysis. Helix propensity was calculated as in Fig. 2A. **(B)** (Left) Contact frequency between L3 segment residues and membrane components for simulations described in (A). Asterisks mark mutated residues. (Right) Additionally, two simulations each were performed for each mutation with the Mid1 C2-PH (residues 580-920) for 1500 ns. Contact frequency analysis was performed on the last 1000 ns except for two simulations: one 4A-NM simulation which never bound and one NM simulation which did not bind until late into the simulation. No contacts were calculated for the unbound 4A-NM simulation. Contacts were calculated for the last 400 ns for the late binding NM simulation in which it was bound. All contact frequency plots are normalized to 1, with the remaining contacts being water (not shown). **(C)** Time sequence of the distance between the L3 and membrane COM along the membrane normal for the indicated Mid1 C2-PH simulations as in Fig. 3B. Data shown is a running average over a 100 ns window.

We additionally performed two simulations for each mutation of the whole C2-PH in two different rotational orientations started apart from the membrane for a duration of 1500 ns (Table 1). The 4A simulations quickly bound the membrane through the L3 loop on timescales similar to that of WT simulations (Figure 5C). One of the NM simulations took longer than 1000 ns to bind through the L3 loop, while the other made several membrane approaches before binding (Figure 5C). While one 4A-NM simulation bound on timescales similar to WT, the other failed to bind during the entire 1500 ns duration (Figure 5C). The membrane contacts within the 677-701 region of the 4A simulations appear less robust compared to WT, though additional contacts are formed with greater frequency in the 658-663 region of the L3 loop (Figure 5B right, compare to Figure 3C). Both the NM and 4A-NM mutations formed drastically fewer contacts with less frequency in comparison to WT (Figure 5B right, compare to Figure 3C). Interestingly, during the NM and 4A-NM simulations where the L3 loop remained unbound for an extended duration (>1 μs), the PH domain also failed to bind, further ruling out PH binding on the timescales available during these simulations.

The results of Figure 5 show that mutation of the NLS sequence (NM) or of both NLS and candidate helix (4A-NM) slow down and weaken association to the studied plasma membrane, in agreement with experiments. However, we did not observe statistically significant effects of 4A mutations. The latter mutation may influence membrane insertion of this region, a process occurring on much longer timescales to our simulations. In all cases, the L3 loop can make robust contacts on short timescales such that the studied mutations do not outright prevent membrane association.

## Discussion

Our results show that the Mid1 monomer has many modes of membrane binding, which may reflect its multiple interactions with membranes and other cytokinetic ring proteins. Figure 6 shows a model of Mid1 in nodes, which takes into consideration super-resolution data of node proteins [21] (slightly shifted by 10 nm to place Mid1 adjacent to the plasma membrane), as well as the estimated size of Mid1’s IDR region based on coarse-grained MD simulations using methods similar to those in Bhattacharjee et al. [38] (data not shown). This figure indicates how a membrane layer of ~8-10 Mid1 per node [21, 39] provides a scaffolding layer for node proteins, together with about twice-as-many membrane-bound Cdc15 and Rng2 [38], as well as Cdr2 (not shown), thus helping anchor formin Cdc12 and type II myosin Myo2. The multiple modes of Mid1 membrane binding may reflect its ability to sustain mechanical forces as Myo2 pulls on Cdc12-nucleated actin filaments, leading to the condensation of the band of nodes into a ring through the opposing steric hindrance by membrane-associated ER [40]. These membrane attachments should also control the resistance of node movement to applied force, an important biophysical parameter for cytokinetic ring organization. Recent evidence suggests that IDR regions of Cdc15 promote condensation of Cdc15 through forces related to liquid-liquid phase separation [38]. Flexibility in Mid1 membrane binding may thus also be related to its ability to participate into a disordered condensate through its IDR region, which is of comparable size to Cdc15.

**Figure 6:**
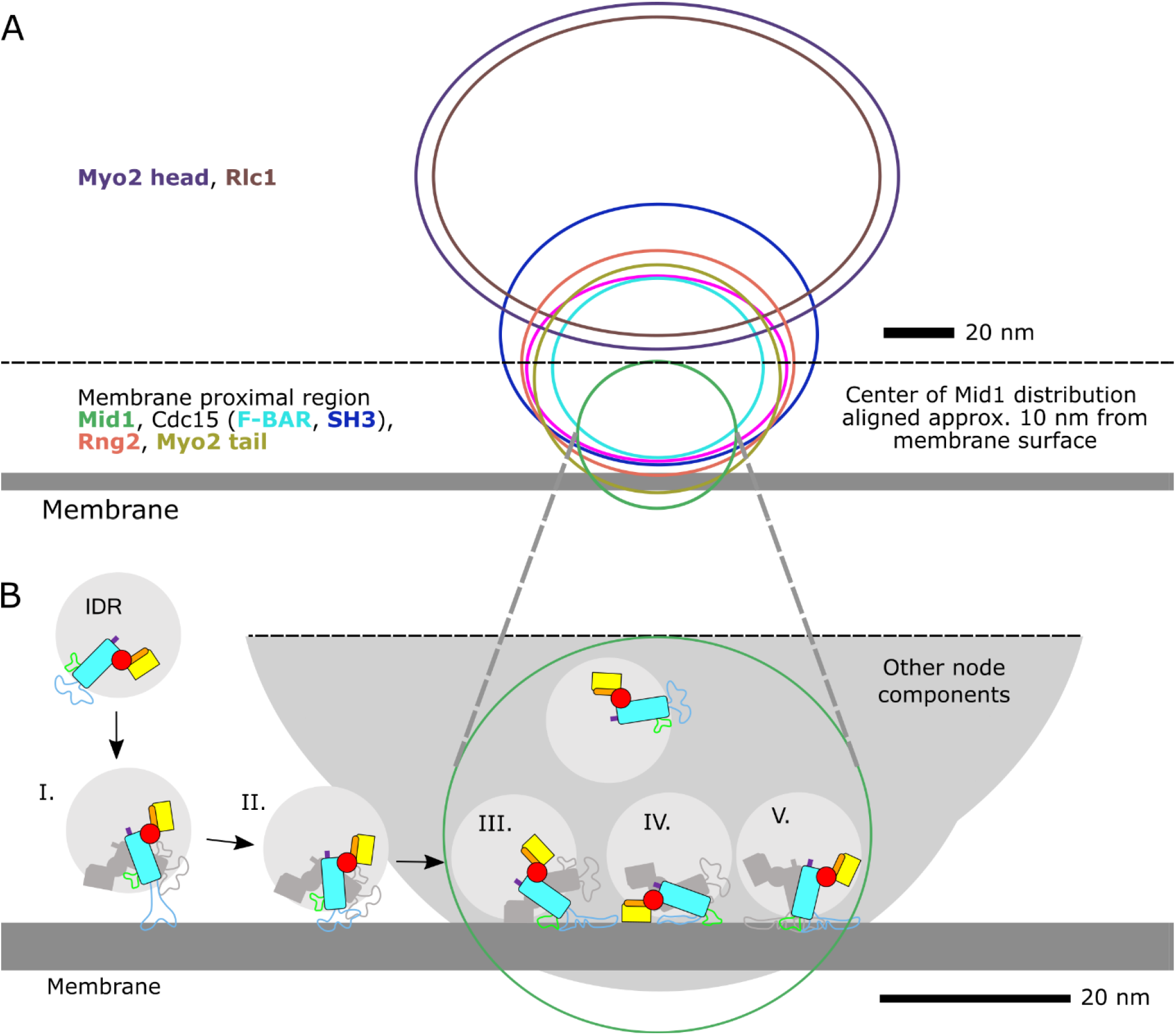
Proposed Mid1 Binding to Cytokinetic Nodes. **(A)** Side view distribution of cytokinetic node components near the membrane as in [21]. Membrane shown 5 nm thick. Distribution of node components has been vertically shifted such that the center of the Mid1 distribution is approximately 10 nm from the surface of the membrane; this is the approximate distance of the connector region from the membrane when the L3 loop binds in an extended way (see (B) I). **(B)** Zoom in on membrane proximal region in (A). Mid1, surrounded by the N-terminal IDR (attachment point in purple) binds initially through the L3 loop and can dimerize using its C2 interface (I). Collapsing further onto the membrane either as a monomer or dimer, Mid1 can form additional contacts in its L3 loop (II), L0 loop (III), or PH domain (IV). Alignment to the Mid1 dimer (Supplemental Figure 2) indicates these configurations may be unfavorable when Mid1 dimerizes. We hypothesize a dimer configuration where the L0 & L3 loop of both Mid1 monomers binds the membrane, in agreement with [9] (V).

In our simulations, the Mid1 monomer initially binds through the L3 loop (Figure 6B.I, B.II), but can further bind through the L0 loop (Figure 6B.III, B.V) and PH domain (Figure 6B.IV). The binding we observe through the L3 loop is fast and primarily driven by positively charged residues making contacts with negatively charge PI lipids, particularly in the NLS sequence and with PIP_2_. However, mutation of the NLS sequence alone is not sufficient to prevent L3 loop association to the membrane in our simulations, with neighboring regions still able to make contacts. The L0 loop contacts, which we observed less often in freely diffusing C2-PH simulations, but appear stable after formation, include hydrophobic residues that insert into the membrane. While we could not observe the PH domain contacting the membrane in freely diffusing C2-PH simulations, the PH domain remains stably bound when started in contact. Residues in the candidate helix region of the L3 loop made membrane contacts, though these were less noticeable compared to the NLS region; we also did not observe stable helix formation, and the hydrophobic residues of the candidate helix were not seen to substantially insert into the membrane. Correspondingly, we did not see a large effect from mutations to the candidate helix, which have been shown to cause Mid1 to be largely cytoplasmic in experiment [25]. It is very likely that helix formation and membrane insertion of hydrophobic or aromatic residues occurs over timescales beyond the reach of serial MD simulations, as has been the case for simulations with liquid-ordered membranes [41]. Overall, these results may indicate that short timescale binding of the L3 loop, in particular the NLS sequence, drives the Mid1 protein’s affinity for PIP_2_ lipids near the center of fission yeast, but that long timescale binding requires making more stable interactions such as those through the L0 loop, PH domain, and candidate helix membrane insertion. This may indicate that the NLS sequence provides an additional signaling role for Mid1 beyond its role in nuclear shuttling, in agreement with studies of the NLS sequence in anillin.

Another way Mid1 strengthens its membrane affinity is through dimerization. While we did not perform simulations of the Mid1 dimer, we wished to see how the Mid1 monomer binding modes we observed may affect dimerization, in particular with regards to problems with steric hinderance and location of membrane binding interfaces. We aligned the dimer structure [9] (Supplemental Figure S2A) using the amino acids involved in the dimer interface of one of the monomers (Supplemental Figure S2B) with the simulated monomer in simulation snapshots, in order to visualize a potential binding partner (Supplemental Figure S2C-F). When the Mid1 monomer is bound solely through the L3 loop, the PH domain of the partner monomer also appears distal to the membrane; however, both the L3 and L0 partner loop would be pointed towards the membrane (Supplemental Figure S2C, D). When the L0 loop is also bound and the C2 domain is quite tilted with respect to the membrane normal, it may be possible for the partner Mid1 monomer to bind its PH domain to the membrane, but this would require some reorientation and/or flexibility of the CNCT region to do so. This also would point the partner Mid1’s L3 and L0 loop away from the membrane (Supplemental Figure S2E). Therefore, we think it is likely simultaneous binding of the L3 loops of both monomers in the dimer would prevent this highly tilted C2 configuration, and therefore the binding of the PH domain; however, the L0 loop would not be prevented from binding in this orientation. When the L3 loop, L0 loop, and PH domain are all bound, the C2 dimerization interface is pointed upwards, and the partner monomer has no steric conflicts (Supplemental Figure S2F). In this orientation, the partner monomer’s PH domain is pointed away from the membrane; however, given the length of the L3 loop, it is possible it could reach the membrane in this orientation.

Taken together, our results suggest that the stable configuration of the Mid1 dimer is simultaneous binding of the L3 and L0 loops of both monomers to the membrane, but not the binding of the PH domain (Figure 6B.V). The PH domain may serve to strengthen the binding of the Mid1 monomer to the membrane, as its binding does not appear to interfere with the availability of the C2 dimerization interface. After dimerization in this manner, the L3 loop could aid in ‘standing up’ of the Mid1 dimer into the stable configuration, causing the detachment of the PH domain.

This begs the question as to the role of the PH domains distal to the membrane. Although binding of PH domains to PI lipids is well studied, PH domains also mediate protein-protein interactions [42, 43]. For example, human anillin, drosophila anillin, and Mid2 PH are known to recruit septins [44–47]. However, little is known about Mid1 PH’s possible role in protein-protein interactions, but it has been suggested it may directly or indirectly interact with ESCRT-associated protein Vps4 [48]. The Mid1 PH domain’s preference for negatively charged PI lipids may indicate a potential role in regulation by phosphorylation, which adds a negatively charged phosphate group to an amino acid. Mid1 is known to be heavily phosphorylated on its N-terminal IDR [7, 34], and regulation by phosphorylation is a common feature of other node proteins [49, 50]. Similar to Cdc15 [38], progressive phosphorylation of the Mid1 IDR may affect the IDR’s association to Mid1’s globular domains. Negatively charged phosphorylated residues could drive the localization of the IDR away from the membrane binding and dimerization interfaces of the C2 domain and towards the PH domain. However, what potential role the PH domain would play in regulation via phosphorylation has not been explored.

In conclusion, this study highlights the multiple binding modes available to monomeric Mid1, with the implications of the results including Mid1 dimerization binding modes, Mid1 localization in the cell, Mid1 regulation, and interfaces available to Mid1 binding partners. Even though our results focused on fission yeast Mid1, they should be relevant for anillin family members more broadly. Fission yeast Mid2, which also plays a stabilizing role for the contractile ring, is predicted by AlphaFold to have a similar geometrical arrangement of C2-PH to that of Mid1’s C2-PH, with most obvious differences in the CNCT, L3, L0, and PH lipid binding regions (compare Supplemental Figure S1C to E). AlphaFold structures of Mid1 and Mid2 of the related medially-dividing *S. japonicus* bear close resemblance to the corresponding structures of *S. pombe* (compare Supplemental Figure S1C to D, E to F). Apart from the extra Rho binding region adjacent to C2, AlphaFold predicted structures of drosophila and human anillin share a similar structure and apparent membrane binding interfaces to those of Mid1 (compare Supplemental Figure S1A, B to C).

## Acknowledgements

We thank Jian-Qiu Wu, Aurelia Honerkamp-Smith, and Thomas Pollard for feedback and suggestions. This work was supported by NIH grant R35GM136372 to D.V. and by NSF grant MCB-2111728 to W.I. This research used resources of the Argonne Leadership Computing Facility, which is a DOE Office of Science User Facility supported under Contract DE-AC02-06CH11357. Portions of this research were conducted on Lehigh University’s Research Computing infrastructure partially supported by NSF Award 2019035 and the high-performance computing capabilities of the Extreme Science and Engineering Discovery Environment (XSEDE), supported by the National Science Foundation, project no. TG-MCB180021.

## Methods

We make use of the crystal structure of Mid1, PDB: 4XOH [9] using the ‘A’ structure and filling in all residues missing in sequence 580-920 using I-TASSER [35] (C2-PH). For simulations of only a section of C2-PH, we simply removed the unneeded residues lines from the filled-in PDB file. Lipid models selected for PIP_2_ included half protonated on the P4 and half on the P5 (CHARMM-GUI models POPI24 & POPI25). Other membrane component models were chosen as named in Fig. 1C. Number of membrane components were chosen to match molar ratios listed in Fig. 1C. L3 segment and PH domain simulations had an initial box size of approximately ~89 Å × 89 Å × 159 Å and ~128,000 atoms, while unbound C2-PH simulations had an initial box size of approximately ~145 Å × 145 Å × 200 Å and ~418,000 atoms to allow protein segments to be placed far enough from the membrane to reorient before binding (Table 1). Additionally, for unbound C2-PH, the protein was started in a different rotational orientation for each independent simulation. C2-PH simulations started on the membrane had a box size of ~138 Å × 138 Å × 143 Å and ~281,000 atoms. (Supplemental Figure S3).

All systems were prepared using CHARMM-GUI [51] *Membrane Builder* [52–55] exporting inputs for GROMACS [56, 57]. Simulations were performed using the CHARMM36(m) force field [56, 58] in GROMACS versions 2018.5 and 2020.4 [59] using the TIP3P water model [60] with 100 mM KCl. Equilibration and production simulations were performed largely as prescribed by the GROMACS inputs generated by CHARMM-GUI. Relaxation included an energy minimization step of 5000 steps using the steepest decent algorithm. Following this, the system was simulated in the NVT (constant particle number, volume, and temperature) ensemble for 250 ps with a 1 fs time step. Then the system was switched to an NPT (constant particle number, pressure, and temperature) ensemble with 1 fs time step for 125 ps, and then a 2 fs time step for 1.5 ns. During equilibration, positional and dihedral restraints were used with gradually decreasing force constants. The duration of equilibration steps was increased for individual cases if needed. Production simulations were performed unrestrained in the NPT ensemble with a 4 fs time step using hydrogen mass repartitioning [61]. Temperature was maintained at 300 K with a Nosé-Hoover thermostat with a time constant of 1 ps. Pressure was maintained at 1 bar with a semi-isotropic Parrinello-Rahman barostat with compressibility of 4.5 × 10^-5^ bar^-1^ and time constant of 5 ps for the unbound C2-PH case, which was increased for all other simulations to 12 ps to improve stability.

Systems were visualized in VMD [62]. Contact analysis was done in python using the *compute_neighbors* function of the MDTraj package [63]. Helix propensity was calculated using DSSP [64, 65] using the GROMACS command *do_dssp* and grouping helixes of the G, H, and I designations (3, 4, and 5 turn helixes respectively). Orientation analysis was done in python using the first principal component as calculated by the *principal* command in GROMACS.

## Supplemental Figures

**Supplemental Figure 1:**
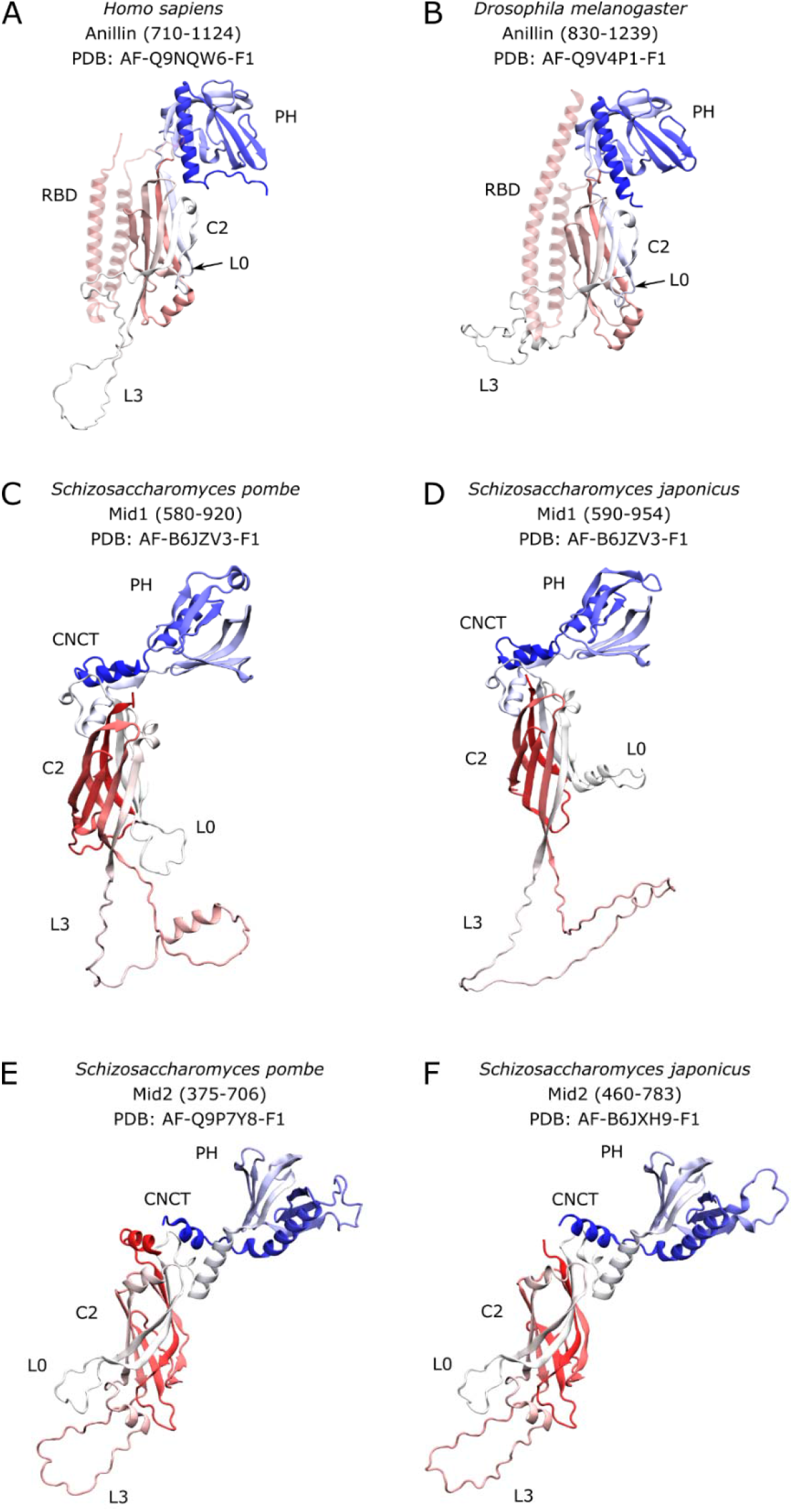
Comparison of AlphaFold structures of anillin and anillin-like Mid1 and Mid2. **(A-F)** Structures are colored with a red to blue gradient from the N to C terminus. The ß7-ß8 loop of the C2 domain is labeled as L0. **(A, B)** The Rho binding domain (RBD) is shown partially transparent to more easily compare Anillin to Mid1 and Mid2.

**Supplemental Figure 2.**
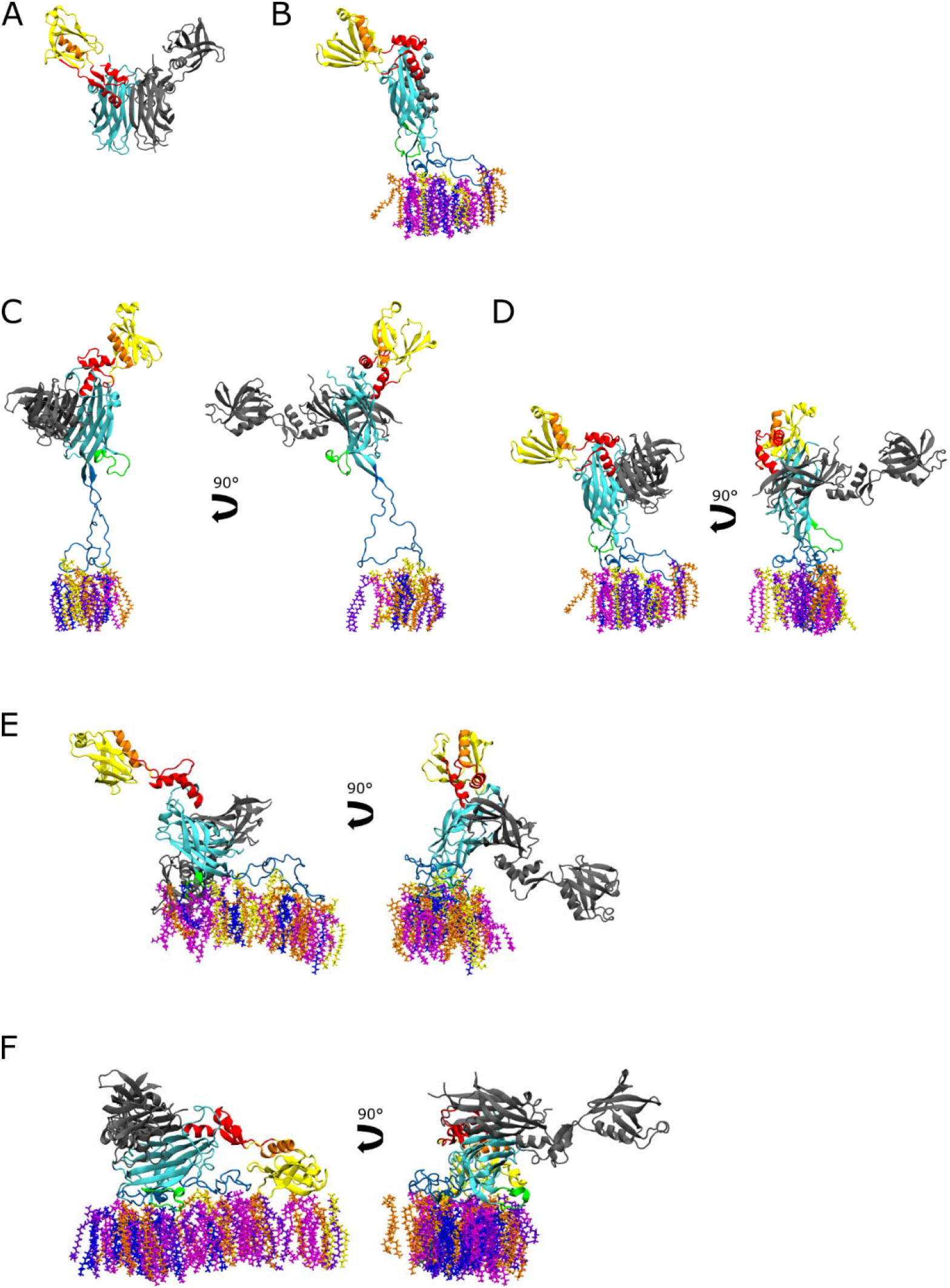
Alignment of Mid1 dimer to monomer bound configurations. **(A)** Dimer structure in PDB: 4XOH. Monomer used for alignment shown in colors matching main figures. Partner monomer shown in grey. **(B)** Mid1 dimerization interfacial residues (spheres); defined as residues of Mid1 monomer which make contact during dimerization (are within 5 Å of the other monomer in PDB 4XOH structure.) **(C-F)** We aligned the residues in the dimerization interface defined in (B) with those in the Mid1 dimer in A. We then visualize the monomer from the all-atom simulation with the partner monomer from alignment (grey). **(C-E)** Binding configurations from Fig. 3A visualized after alignment. **(F)** Final binding configuration from Fig. 4A visualized after alignment.

**Supplemental Figure 3:**
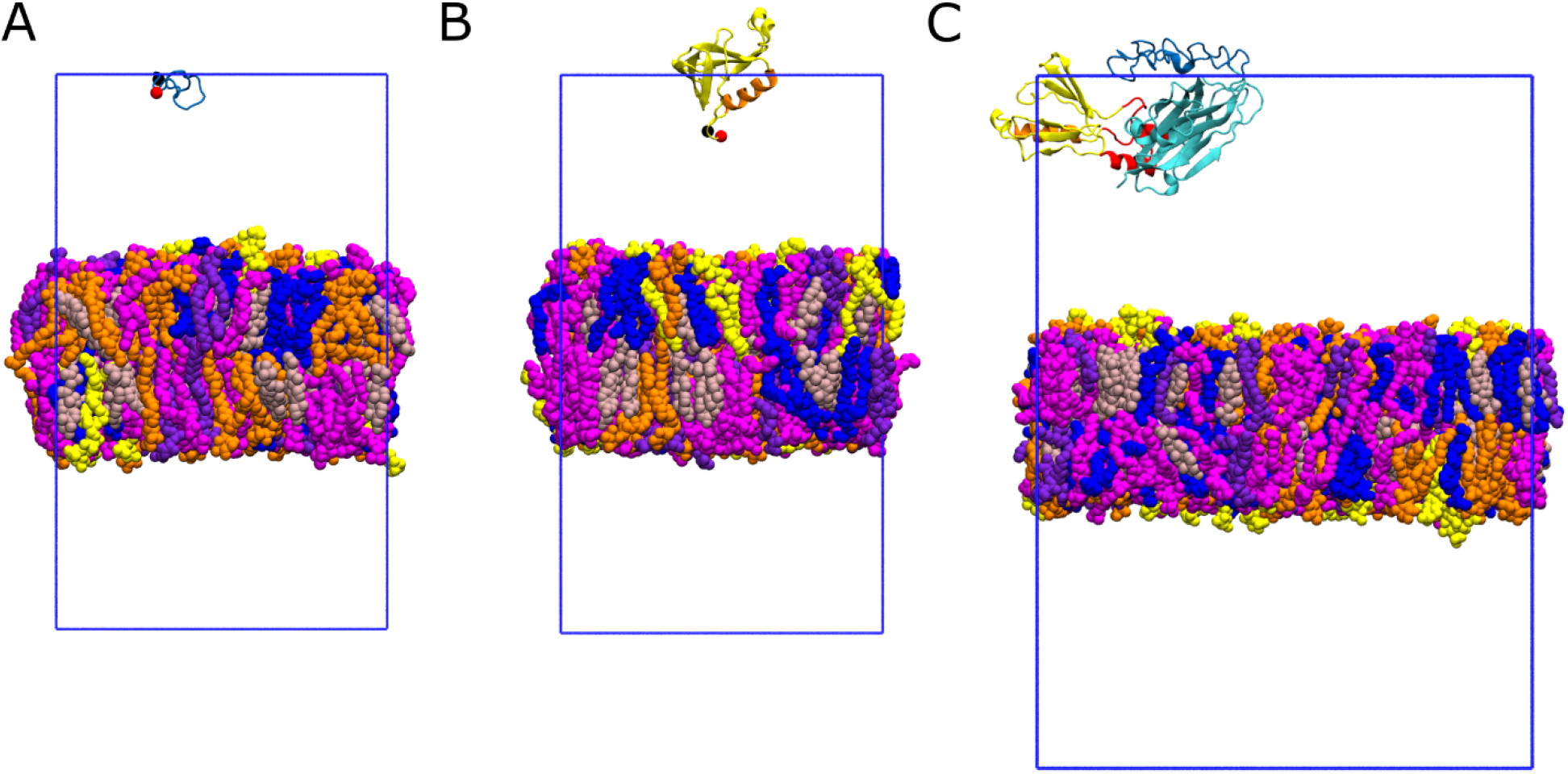
Simulation Initial Conditions. **(A-C)** Initial condition of simulations after equilibration described in Methods. Blue rectangle indicates the periodic boundary. Molecules are rendered across periodic boundaries via GROMACS operation *trjconv* with flag ‘-*pbc whole’*. **(A)** Initial condition snapshot of L3 segment simulations. **(B)** Initial condition snapshot of PH domain simulations. **(C)** Initial condition snapshot of a freely diffusing C2-PH simulation. Each independent simulation used a different rotational orientation; one example is shown.

## Movies

**Movie 1: Simulation of L3 segment.** Representative simulation from those described in Figure 2A, B

**Movie 2: Simulation of PH domain.** Representative simulation from those described in Figure 2C, D

**Movie 3: Simulation of C2-PH unbound.** Representative simulation from those described in Figure 3

**Movie 4: Simulation of C2 PH unbound with L0 insertion.** Simulation from those described in Figure 3 in which the L0 loop inserts into the membrane.

**Movie 5 Simulation of C2-PH bound.** Representative simulation from those described in Figure 4

**Movie 6: Simulation of L3 segment 4A.** Representative simulation of the L3 segment with the 4A mutation from those described in Figure 5A and B left.

**Movie 7: Simulation of L3 segment NM.** Representative simulation of the L3 segment with the NM mutation from those described in Figure 5A and B left.

**Movie 8: Simulation of L3 segment 4A-NM.** Representative simulation of the L3 segment with the 4A-NM mutation from those described in Figure 5A and B left.

